# Democratizing Data-Independent Acquisition Proteomics Analysis on Public Cloud Infrastructures Via The Galaxy Framework

**DOI:** 10.1101/2021.07.21.453197

**Authors:** Matthias Fahrner, Melanie Christine Föll, Björn Grüning, Matthias Bernt, Hannes Röst, Oliver Schilling

## Abstract

Data-independent acquisition (DIA) has become an important approach in global, mass spectrometric proteomic studies because it provides in-depth insights into the molecular variety of biological systems. However, DIA data analysis remains challenging due to the high complexity and large data and sample size, which require specialized software and large computing infrastructures. Most available open-source DIA software necessitate basic programming skills and cover only a fraction of the analysis steps, often yielding a complex of multiple software tools, severely limiting usability and reproducibility. To overcome this hurdle, we have integrated a suite of DIA tools in the Galaxy framework for reproducible and version-controlled data processing. The DIA suite includes OpenSwath, PyProphet, diapysef and swath2stats. We have compiled functional Galaxy pipelines for DIA processing, which provide a web-based graphical user interface to these pre-installed and pre-configured tools for their usage on freely accessible, powerful computational resources of the Galaxy framework. This approach also enables seamless sharing workflows with full configuration in addition to sharing raw data and results. We demonstrate usability of the all-in-one DIA pipeline in Galaxy by the analysis of a spike-in case study dataset. Additionally, extensive training material is provided, to further increase access for the proteomics community.

## Background

Data-independent acquisition (DIA) is a recently developed method addressing the need for reproducible and robust explorative proteomic measurements in larger sample cohorts ^1^. Compared to classical data-dependent acquisition (DDA) measurements, in DIA all MS1 precursor peptide ions within a predefined m/z range (“window”) are fragmented and subjected to MS2 scans. Especially for high-throughput studies with dozens of samples, DIA has been shown to yield higher numbers of identifications and quantifications ^2–4^. Furthermore, when compared to isobaric labelling approaches, DIA is less susceptible to batch effects stemming from chemical tagging and allows for quantitative proteome comparison in large cohorts ^5,6^. Multiple DIA strategies have been developed over the last decade as reviewed in Ludwig et al. ^3^. Applying overlapping isolation windows and subsequent demultiplexing has been shown to improve the precursor selectivity ^7,8^. However, for overlapping isolation window DIA data, specific data processing may be required to ensure compatibility with subsequent data processing steps.

Different data processing strategies have been developed for DIA data, where the most common strategies apply a spectral library to enable confident identification of peptides in DIA data ^4^. Due to the complex MS2 spectra as well as the requirement for *a priori* knowledge a complete DIA data analysis can be divided into three steps: (i) the generation of a spectral library (ii) the actual identification by matching measured fragment masses and their respective retention times to the precursor and fragment information within the spectral library (iii) a statistical follow-up analysis yielding the identification of significantly altered protein expression profiles. A prototypical DIA analysis often includes a multitude of software and system environments for steps such as spectral library generation, peptide and protein identification in DIA measurements, and differential statistics (**Figure 2A, upper panel**). Spectral libraries are often based on DDA data analyses e.g. using MaxQuant ^9^ via a graphical user interface, followed by library generation e.g. using diapysef ^10,11^ in a python shell, and OpenSwath ^10^ tools on the command line for library refinement. For the analysis of DIA data containing overlapping isolation windows, a demultiplexing step may be required, e.g. during the conversion from vendor-specific file formats to the open mzML ^12^ format with tools such as msconvert ^13^. The identification of peptides in DIA data can be performed using a variety of software suites e.g. OpenSwathWorkflow ^14^ followed by FDR scoring using PyProphet ^15^. The peptide identification and quantification, target decoy scoring as well as the results export can be performed in the standard Windows terminal or an enhanced terminal for Windows such as MobaXterm. The final differential expression analysis can be performed using specialized software such as MSstats in R programming language. This portrayal serves to illustrate the inherent complexity of modular data analysis in modern DIA proteomics.

The multi-step characteristic of a complete DIA data analysis has encouraged the development of a variety of software options, some of which are particularly powerful in one or more of the three steps ^4,16^. However, the usage of multiple software packages impedes streamlined high-throughput analysis and poses hurdles for software compatibility and reproducibility. Hence, DIA data analysis, especially in the context of powerfully adaptable modular software tools, requires an advanced level of computational skills for software installation, connecting them into analysis workflows and usage in the case of command-line-based software. Recent endeavors have used so-called Docker-based structures to distribute pre-assembled and readily usable DIA software bundles, however, they still require a high degree of computational, especially programming skills ^17–19^. Thus, a hidden requirement for DIA data analysis has been sophisticated bioinformatic skills, due to the involvement of multiple software tools in a complete analysis and individual analysis steps, that are performed using open-source programming languages such as R or Python. To enable straightforward and user-friendly analysis, monolithic software such as Spectronaut and Skyline has been developed ^2,7^. However, it remains challenging to embed monolithic software in workflow environments and to enable compatibility and interoperability with other software. Moreover, their design often lacks the ultimate flexibility and tunability of modular software suites such as OpenSwath ^10^.

OpenSwathWorkflow (OSW) is one of the earliest open-source DIA analysis software suites, that supports a large number of functionalities and parameters allowing for a fully customized DIA analysis ^20^. Yet, the sophisticated flexibility and numerous parameter options make it difficult to report all crucial settings e.g. in scientific communications; potentially limiting reproducibility and transparency of DIA analysis. Moreover, OSW by default does not have a graphical user interface, limiting its accessibility and usability to researchers that are familiar with executing software from the command line. The urgent need for a user-friendly and fully customizable DIA analysis pipeline is highlighted by recent endeavors in streamlined DIA analysis options applying OSW ^18,19,21^.

Here we present a user-friendly repertoire of DIA analysis tools, which can be accessed by a broad user community via the web-based analysis and workflow framework Galaxy ^22^. The Galaxy framework makes thousands of bioinformatics tools available to the scientific community without requiring advanced bioinformatic or programming skills. Galaxy analyses are stored in so-called histories, in which all tool names, tool versions, tool parameters and intermediate data are saved, hence representing an important step for reproducibility. More than a hundred public Galaxy servers are available worldwide and offer access to powerful public cloud infrastructure for academic or non-commercial purposes. Into this powerful framework, we have integrated a suite of eleven modular DIA tools based on OpenSWATH ^10^, diapysef ^11^, PyProphet ^15^, and swath2stats ^23^ (**Table 1**). Together with existing Galaxy tools all necessary DIA analysis steps can be executed within Galaxy with high flexibility and in an easily accessible manner. We apply the DIA analysis tools on an *E*.*coli*:HEK spike-in dataset to demonstrate the use of a Galaxy-based DIA Analysis pipeline that facilitates standardization and reproducibility and is compatible with the principles of FAIR (findable, accessible, interoperable, and re-usable) data and MIAPE (minimum information about a proteomics experiment) ^24,25^.

**Table 1.**
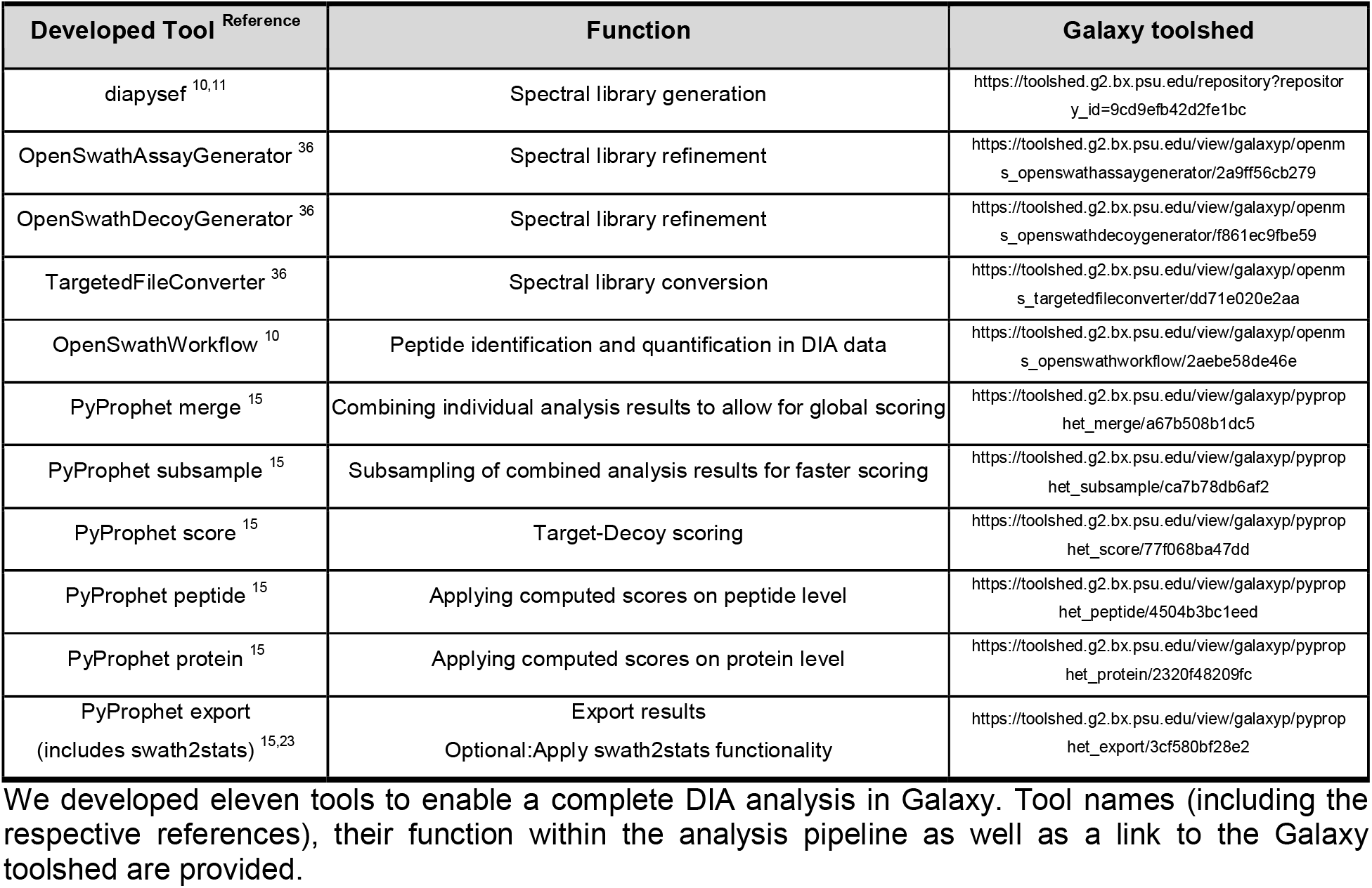
Overview of developed tools for the DIA data analysis in Galaxy

## Methods

### *Escherichia coli* K12 (*E*.*coli*) and human embryonic kidney 293T (HEK) whole proteome samples

*E*.*coli* and HEK proteome samples were prepared as previously described ^26^. Briefly, cells were lysed using 5% SDS in 50 mM triethylammonium bicarbonate (TEAB) at pH 7.55 by applying sonication (20 cycles with 30/30 sec on/off high energy) with a Bioruptor device (Diagenode, Liège, Belgium). Following centrifugation for 8 min at 13,000 g, proteins in the supernatant were reduced by incubating with 5 mM TCEP (Sigma) at 95°C for 10 min and subsequently alkylated by incubating with 5 mM IAA at room temperature in the dark. Protein digestion and purification was performed on S-Trap™ micro spin columns (Protifi, Huntington, NY) according to the manufacturer’s protocol. After elution the peptide concentrations were measured using a bicinchoninic acid assay (Thermo Scientific) according to the manufacturer’s protocol. Different amounts of *E*.*coli* peptides (0, 0.05, 0.15, 0.4 and 0.8 µg) were added to stable amounts of HEK peptides (2.5 µg) resulting in the following ratios: HEK only; 1:50; 1:17; 1:7 and 1:3. Two replicates of each *E*.*coli*:HEK ratio were prepared. Samples were vacuum-concentrated until dryness and stored at -80°C until LC-MS/MS analysis.

### LC-MS/MS analysis

One ug of peptides were analyzed on a Q-Exactive Plus mass spectrometer (Thermo Scientific, San Jose, CA) coupled to an EASY-nLC™ 1000 UHPLC system (Thermo Scientific). The analytical column was self-packed with silica beads coated with C18 (Reprosil Pur C18-AQ, d = 3 Â) (Dr. Maisch HPLC GmbH, Ammerbusch, Germany). For peptide separation, a two-step linear gradient with increasing amount of buffer B (0.1% formic acid in 80% acetonitrile, Fluka) was applied, ranging from 8 to 43% buffer B over 90 min and from 43 to 65% buffer B in the subsequent 20 min (110 min separating gradient length). Additionally, buffer A and buffer B contained 3% ethylene glycol (final concentration), which has been shown to improve electrospray ionization ^27^. For the spectral library one representative sample of each *E*.*coli*:HEK ratio (in total n = 5 samples) was measured using data-dependent acquisition. Briefly, survey scans covering an m/z range from 385 to 1015 m/z were performed at 70,000 resolution, an AGC target of 3e6 and a maximum injection time of 50 ms followed by targeting the top 10 precursor ions for fragmentation scans at 17,500 resolution with 1.6 m/z isolation windows, a stepped NCE of 25 and 30, and an dynamic exclusion time of 35 s. For all MS2 scans the intensity threshold was set to 6.3e4, the AGC to 1e5 and the maximum injection time to 160 ms. The twenty *E*.*coli*:HEK ratios samples were measured using data-independent acquisition. For DIA two cycles of 24 m/z broad windows ranging from 400 to 1000 m/z with a 50% shift between the cycles (staggered window schema was used) ^28^. MS2 scans were performed at 17,500 resolution, an AGC target of 1e5 and a maximum injection time of 80 ms using stepped NCE of 25 and 30. After 25 consecutive MS2 scans a MS1 survey scan was triggered covering the same range and with the same settings as in the DDA measurements.

### Data analysis using Galaxy

The complete data analysis was performed on the European Galaxy server ^22^. The analysis history for the spectral library generation and the DIA analysis (including the statistical analysis) have been published via Galaxy ^29,30^ and can be found in the additional data. Briefly, spectral library generation was performed by analyzing five DDA measurements representing different *E*.*coli*:HEK ratios using MaxQuant in Galaxy. A reviewed human protein database containing 20,426 sequences (08/06/2019) and an *E*.*coli* protein database containing 4,352 sequences (03/28/2019) were retrieved from UniProt. The five DDA measurements were specified as fractions to yield a combined peptide and protein identification. For peptide identification, fully tryptic digestion (Trypsin/P) was assumed allowing for up to one missed cleavage and at least one unique peptide per protein was requested. Carbamidomethylation(C) of cysteine was set as a fixed modification, whereas oxidation(M) on methionine was applied as variable modification. Search results were filtered for 1 % FDR on both, peptide spectrum match (PSM) as well as protein level. Unique identified peptides, as well as a list of reference peptides (iRT peptides), were used to generate a spectral library with diapysef. The retention time alignment method was set to linear. The spectral library was refined using OpenSwathAssayGenerator with the default settings except for a more stringent m/z threshold of 0.015 Thompson for both, the precursor ion selection as well as the fragment ion annotation. Furthermore, a mass range between 400 and 1000 Thompson for precursor ions was considered. To allow for subsequent FDR scoring, shuffled decoy transitions were added using the OpenSwathDecoyGenerator and setting the m/z threshold to 0.015 Thompson for the fragment ion annotation. In a final step, the spectral library was converted from a tab-separated values (tsv) file format to the peptide query parameter (pqp) format using the TargetedFileConverter. Peptide Identification of the DIA measurements was performed using the freshly built spectral library in combination with the same list of reference peptides (iRT peptides) that was already used during the library generation. For the DIA analysis, OpenSwathWorkflow with default settings and a few adjustments was used. The m/z extraction window was set to 20 ppm on MS/MS and 10 ppm on MS1 level. Within the “Parameters for the RTNormalization for iRT peptides” section the “outlier detection method” was set to none and “choose the best peptides based on their peak shape for normalization” was enabled. A minimal number of 7 iRT peptides was requested and 20 ppm mass tolerance for the iRT transitions was applied. In the “Scoring parameters” section the minimal peak width was set to 5.0 and the computation of a quality value was enabled. The usage of mutual information (MI) scores was deactivated. The “Use the retention time normalization peptide MS2 masses to perform a mass correction” was set to regression_delta_ppm. OpenSwathWorkflow results of each DIA measurement were combined into a single file using the PyProphet merge tool. Target-decoy scoring of the merged OpenSwathWorkflow results was performed using XGBoost as a classifier in the PyProphet score tool. Computed target-decoy scores were applied on peptide and protein level in an experiment-wide and a global context to estimate protein-level FDR control using the PyProphet peptide and PyProphet protein tool, respectively. DIA analysis results were exported as a tsv file using the PyProphet export tool. Since peptide and protein inference in the global context was conducted, the exported results were filtered to 1% FDR by default. Additionally, the swath2stats functionality was used to provide a summary file, a protein and peptide signal table as well as a MSstats input tsv file. Statistical analysis was performed using the MSstats tool as well as a comparison annotation file in two different ways: (i) using the MSstats input tsv file generated using the swath2stats functionality and (ii) using the PyProphet export tsv file and an MSstats sample annotation file. For the two approaches, the “input source” parameter needs to be adjusted and set to “MSstats 10 column format” when using the swath2stats prepared tsv file or “OpenSWATH” when using the PyProphet tsv file.

## Findings

### Galaxy enables easily accessible, straightforward and reproducible DIA data analysis

Here we present an all-in-one DIA analysis pipeline within the Galaxy framework, enabling easy access to a suite of advanced software tools and providing sufficient computational resources for large-scale DIA data analyses. We developed and implemented all necessary tools, required for a complete DIA data analysis into the Galaxy framework (**Figure 1A and Table 1**). The newly developed tools were integrated with state of the art proteomic Galaxy tools such as MaxQuant, MSstats and basic text manipulation tools to build a functional DIA analysis pipeline. All newly integrated DIA tools are based on open-source software such as diapysef, OpenSwath, PyProphet, swath2stats ^10,11,23^. The tools were built in a modular way, allowing a fully customized analysis. Each analysis step can be executed individually or assembled as a workflow in the Galaxy platform facilitating a streamlined and straightforward analysis (**Figure 1A**) ^14,20,31^. All parameter options of the original software can be modified via the Galaxy GUI, providing a maximum of user-adjustable configurations for fine-tuning of analysis. We consider archiving of such details to be relevant for reproducibility; deposition and sharing of entire Galaxy workflows is a very straightforward and integrated way of doing so. In fact, published Galaxy histories include complete provenance data, allowing to reproduce the same analysis that has been published. Thus, version-control and version-archiving constitute a major feature of this approach.

**Figure 1.**
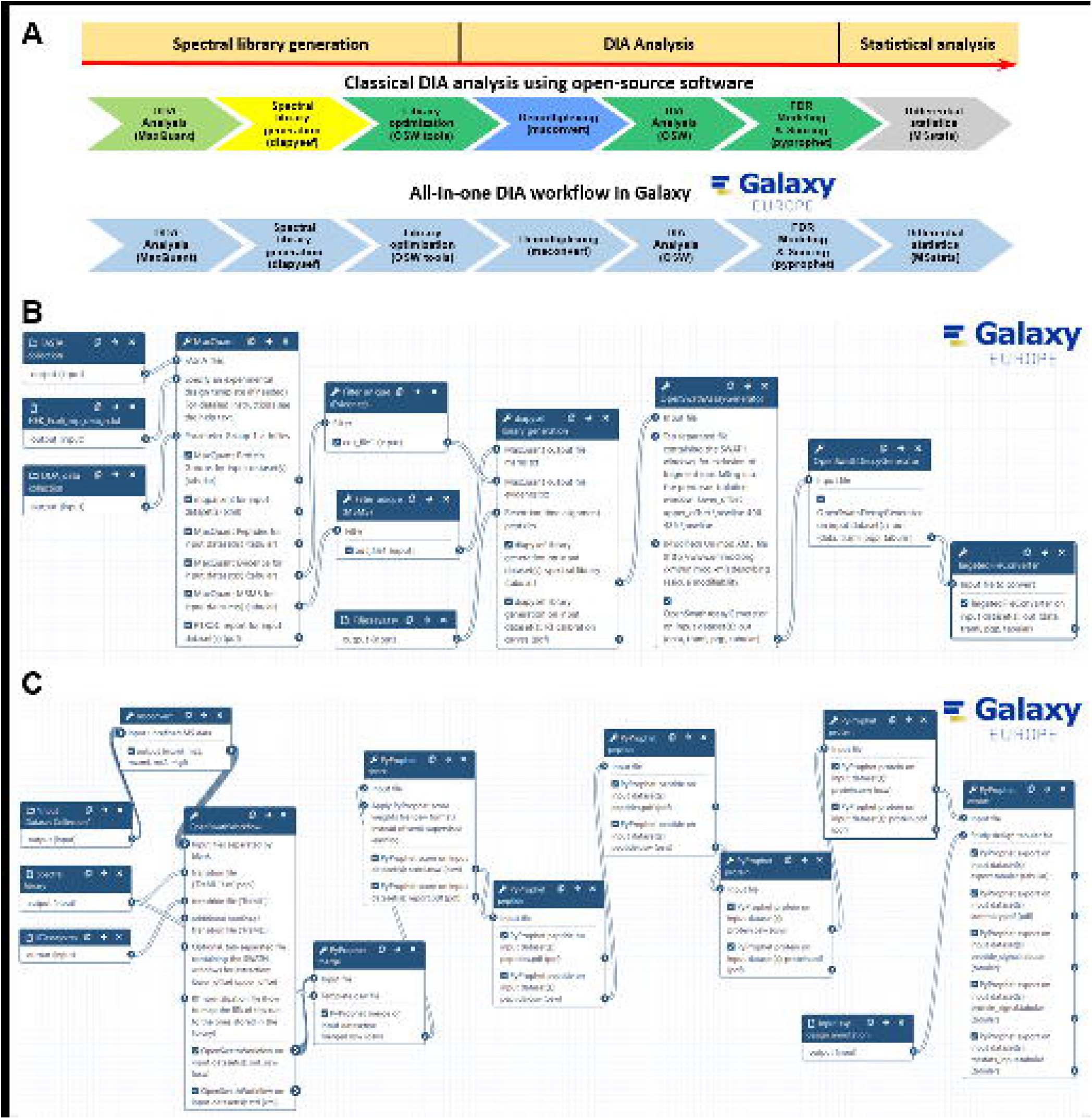
Introducing an All-in-one DIA analysis solution by implementing all necessary tools for a DIA analysis into the Galaxy framework. (A) Schematic overview of an classical data-independent acquisition (DIA) analysis workflow as compared to the here introduced All-in-one workflow in the Galaxy framework. The classic DIA workflow includes different software environments and operating system requirements as indicated by different color (light green: local MaxQuant analysis; yellow: diapysef python shell; dark green: MobaXterm enhanced terminal for Windows; blue: local msconvert; grey: MSstats in Rstudio), whereas all necessary tools are now implemented into the Galaxy framework. A complete DIA analysis can be divided into three steps: (i) Spectral library generation (ii) Peptide and protein identification and quantification in DIA data (iii) Statistical analysis to identify differentially expressed proteins. (B) Generation of a spectral library based on the analysis of data-dependent acquisition (DDA) analysis shown as Galaxy workflow. (C) DIA data analysis shown as Galaxy workflow.

**Figure 2.**
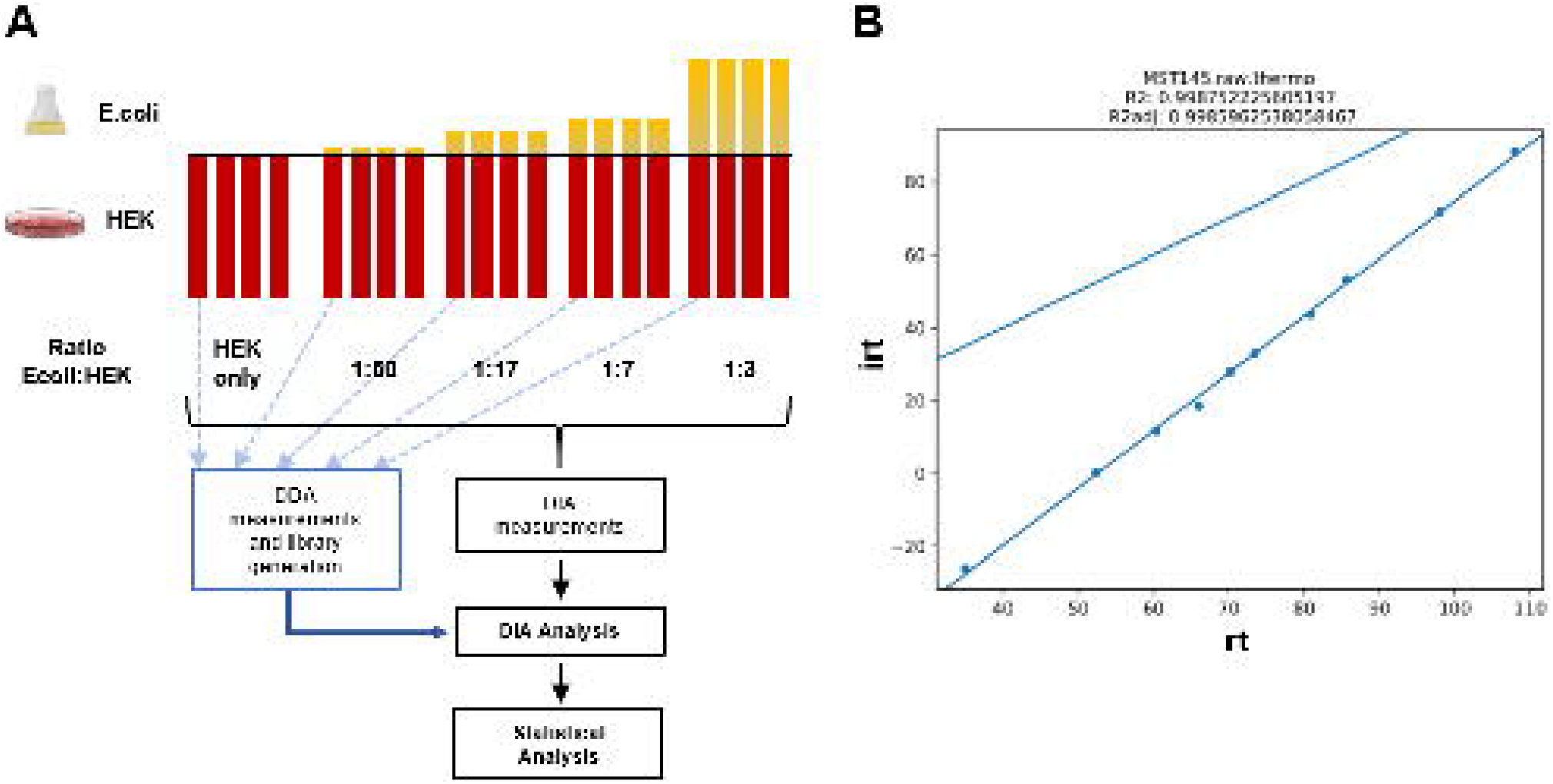
Analysis of a DIA spike-in dataset in Galaxy. (A) Experimental design of a spike-in dataset based on equal amounts of HEK proteome and known spike-in amounts of E.coli proteome. For spectral library generation one representative sample of each mixture was measured using data-dependent acquisition (DDA). Each E.coli:HEK ratio was measured in four replicates using data-independent acquisition (DIA). DIA analysis was performed based on the spectral library and the individual DIA measurements followed by statistical analysis to identify differentially expressed proteins. (B) Retention time (rt) alignment plot of the measured rt and respective indexed retention time (irt) of reference peptides (iRT peptides) during the generation of the spectral library (exemplarily shown for one of the DDA measurements). All measured rt are converted to irts based on the linear regression of the reference peptides (R^2^ and R^2^ adjusted for the linear regression are shown above the plot).

We present a Galaxy-based, complete DIA analysis pipeline that consists of three major parts: (i) spectral library generation using DDA data (**Figure 1B**) ^32^; (ii) peptide (and protein) identification and quantification in DIA data using the aforementioned spectral library (**Figure 1C**) ^33^; and (iii) statistical analysis (**Supplementary Figure 1**) ^34,35^. The workflows for each step have been published via Galaxy and can be downloaded and adjusted or directly run in Galaxy ^32–35^.

We wish to emphasize that embedding these tools in Galaxy not only enables the user-friendly usage but also fosters a new level of reporting and reproducibility in DIA proteomics: the tools as such may be combined in different ways and many tools provide an array of user-adjustable fine-tuning parameters. For illustration, the basic DIA analysis workflow (provided as a training and discussed in further detail below) includes, among others, the three tools OpenSwathWorkflow, PyProphet score and PyProphet peptide each of which with more than 10 fine-tuning settings. This results in numerous possible parameter-combinations which, to our knowledge, are rarely reported in detail. The Galaxy framework addresses this issue by the possibility of depositing and sharing entire workflows (including complete provenance data), also in the context of scientific publications. We consider this feature to be a major benefit of our approach of integrating a DIA processing suite in Galaxy.

### Democratizing data-independent acquisition analysis

The web-based access and graphical user interface in Galaxy empowers a broad community of researchers to perform DIA data analysis. All DIA Galaxy tools are pre-installed and ready to be used on several public Galaxy servers, for example, the European Galaxy server ^22^. Running on public clouds, these Galaxy servers provide an immense computational power that enables running many DIA analyses in parallel without needing to invest or block private computing resources. Most of the software that we integrated into Galaxy is normally only usable with basic programming skills in R or python and thus excludes many proteomics researchers from using them. With the integration into Galaxy, these software tools are now usable by a much broader community via Galaxy’s graphical user interface that allows to specify all input files and parameters and to build analysis workflows based on modular tools. In the following sections, we present more detailed insight into the various steps of Galaxy-supported DIA analysis using the aforementioned tool suite.

### Spectral library generation based on data-dependent acquisition measurements

A spectral library is generated with the newly developed diapysef tool using either the proteotypic peptides or the unfiltered MaxQuant results in combination with a list of reference peptides, to which the RTs of all identified peptides will be aligned using either a linear or a non-linear regression ^37,38^. The diapysef tool in Galaxy automatically generates a pdf file containing the RT calibration curves, highlighting the identified reference peptides and the respective linear or non-linear regression fit (**Figure 2B**). The calibration curves provide a valuable overview of the suitability of the reference peptide with regards to their linear elution as well as the reproducibility of the identification and elution in the analyzed samples/fractions. To improve the sensitivity and selectivity for the detection of typical peptides the spectral library can be refined using the OpenSwathAssayGenerator tool ^36^. Briefly, the number of transitions per precursor ion is reduced and precursors can be filtered to fit the covered mass-to-charge range of the DIA measurements (typically between 400 - 1000 m/z). To enable false discovery rate (FDR) based scoring, an equal number of decoy transitions can be added to the spectral library using the OpenSwathDecoyGenerator tool. In an optional step, the spectral library can be converted to the required format (traml, tsv or pqp) using the TargetedFileConverter tool. In particular, for the generation of result files in the osw format using OpenSwathWorkflow, the spectral library is required as a pqp file.

### OpenSwath in Galaxy allows for the versatile, reproducible, and robust DIA analysis of large proteomic cohorts ^10^

The peptide identification and quantification of the individual DIA measurements is performed in the OpenSwathWorkflow (OSW) tool in Galaxy using the freshly built spectral library, a list of reference peptides (RT peptides) as well as the demultiplexed DIA files in the open mzML format. In most studies, multiple DIA measurements are performed and the different samples should be compared qualitatively and/or quantitatively. Therefore, the individual OSW result files can be merged using the spectral library as a template in the PyProphet merge tool. The target-decoy scoring is performed by applying semi-supervised learning and an error-rate estimation using the PyProphet score tool on the merged OSW results. Noteworthy, the semi-supervised learning and error rate estimation of a merged file containing several 100s of individual DIA measurements can require considerable computational resources. To decrease the analysis time of the semi-supervised learning, the merged OSW results can be first subsampled using the PyProphet subsample tool and subsequently scored using the PyProphet score tool. The computed scores can be applied to the complete merged OSW results. The PyProphet score tool in Galaxy generates an overview of the sensitivity and specificity of the target-decoy scoring as well as a visualization of the distributions of target and decoy transitions **(Figure 3)**. To conduct peptide and protein inference in run-specific, experiment-wide or global context the tools PyProphet peptide and PyProphet protein can be used, respectively ^39^. Each step will generate an overview of the scores and the resulting target-decoy distributions (**Supplementary Figures 2-5**). Afterwards, peptide identification and quantification results can be exported as a tsv file using the PyProphet export tool in Galaxy. Furthermore, we integrated the swath2stats functionality into the PyProphet export tool, allowing the user to visualize the analysis results and to further process the results, e.g. providing an MSstats compatible input tsv ^23^.

**Figure 3.**
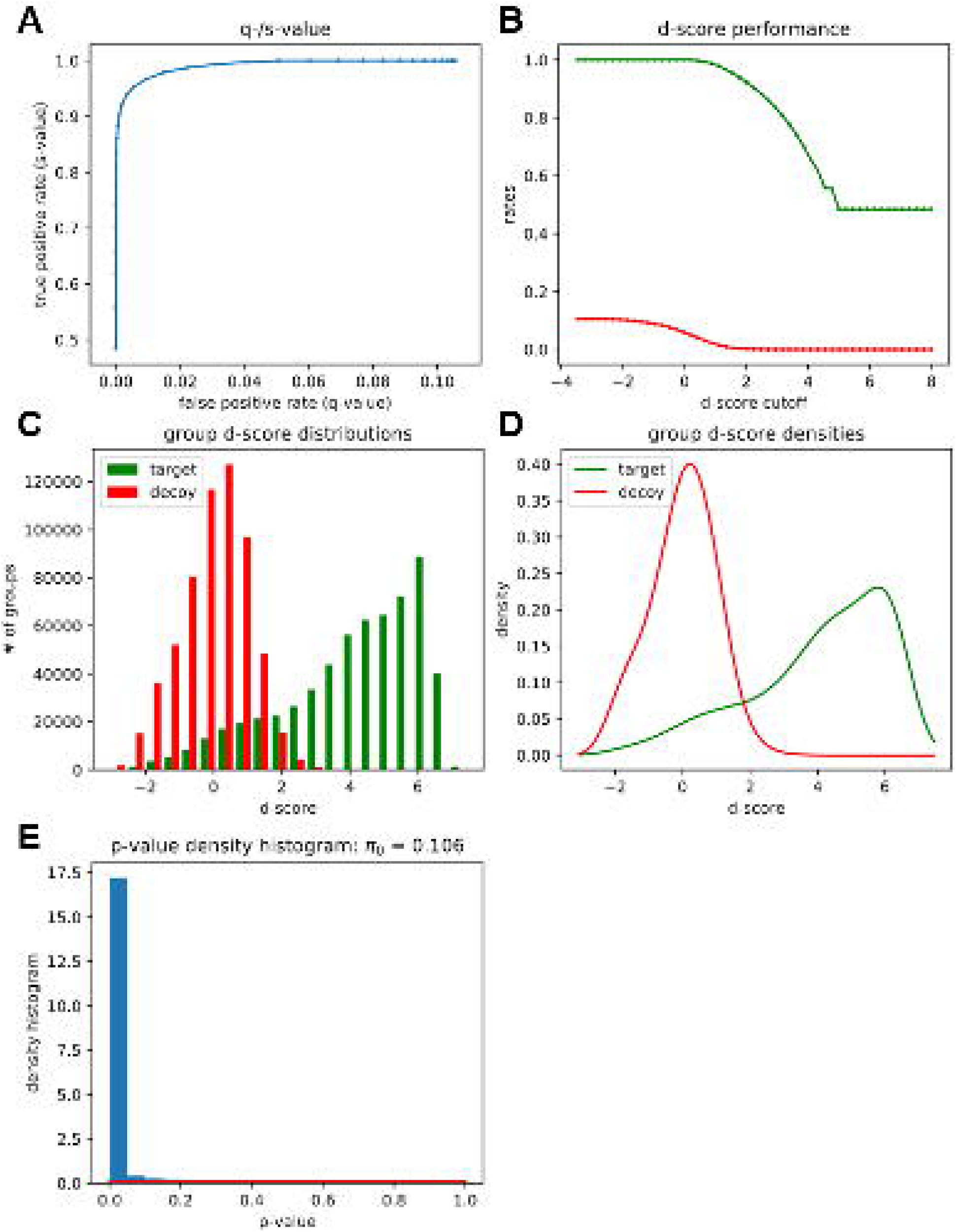
Overview of Target-Decoy scoring using PyProphet score during the DIA analysis in Galaxy. (A) Receiver operating characteristic (ROC) curve highlighting the sensitivity and specificity of the Target-decoy scoring. (B) Blot showing the discriminatory score (d-score) performance between target (green) and decoy (red) transitions. (C) and (D) Barblot and density plot showing d-score distribution among target (green) and decoy (red) transitions. (E) Histogram showing the distribution of p-values computed based on the target-decoy scoring.

### MSstats enables statistical relative quantification of proteins and peptides in DIA proteomics

In most quantitative proteomic studies a statistical analysis is performed to identify significantly altered peptide and protein profiles between different samples. MSstats is a specialized R programming language package for the statistical analysis of proteomic data that has recently been implemented as a Galaxy tool. Differential expression analysis of DIA data can be performed using the swath2stats processed tsv file or by using the non-processed output of the PyProphet export tool in combination with a sample annotation and a contrast matrix file with the MSstats tool.

#### Accessibility and training

Our Galaxy DIA tools are accompanied by hands-on training material, which we have developed and made available via the central Galaxy Training Network ^40,41^. The DIA analysis training material is split into the three major steps: (i) generation of a spectral library ^42^, (ii) DIA data analysis ^43^, and (iii) statistical analysis ^44^. The web addresses of the online trainings are provided as references ^42–44^. Each training consists of a brief introduction to the theory and principles of the respective analysis step. A concise set of training data is provided via a publicly available deposit ^26,45^. Users can directly load the training data into a Galaxy history via the Galaxy URL upload functionality. Using either the training data or their own input data, users can follow the step-by-step introduction provided in the training material. To increase the learning experience through active participation each training includes questions regarding intermediate results based on the provided training data. Of note, intermediate results of rather time-consuming analysis steps are provided, limiting the time consumption of each training. With the extensive set of Galaxy training material for a complete DIA analysis, we wish to enable efficient online self-education of researchers around the world; a topic which has gained increasing interest due to an avalanche of recent, pandemic-related travel restrictions ^46^.

#### Case Study

To illustrate the functionality and utility of our DIA analysis pipeline, we analyzed a DIA dataset, representing a human cell line proteome (human embryonic kidney (HEK) cells) with known spike-in amounts of a distinguishable bacterial *Escherichia coli* (*E*.*coli)* proteome (**Figure 2A**). Additionally, all samples contain a set of 11 synthetic reference peptides for the retention time (RT) alignment ^37^. The dataset includes *E*.*coli*:HEK ratios ranging from 1:50 to 1:3, reflecting a dynamic range of the altered protein abundances. For each *E*.*coli*:HEK ratio n = 4 replicates were measured resulting in a total of 20 DIA measurements. For spectral library generation, one representative sample of each *E*.*coli*:HEK ratio was measured using DDA and analyzed with the MaxQuant tool in Galaxy. The Galaxy framework allows combining all compatible tools. We applied a basic Galaxy text manipulation tool to filter the peptide and protein identifications tsv file for proteotypic peptides, to avoid ambiguous peptides that potentially originate from various proteins. The spiked-in reference peptides elute linearly as highlighted by the RT calibration curves (**Figure 2B**, exemplarily shown for one of the DDA measurements). The staggered window schema of the DIA measurements required demultiplexing before the analysis with OpenSwathWorkflow, which was performed using the msconvert tool in Galaxy. The analysis of the 20 DIA measurements with the OpenSwathWorkflow tool in Galaxy results in the sensitive and selective identification of target transitions (**Figure 3A**). Furthermore, target and decoy transitions show distinct distributions based on the computed d-scores with the PyProphet score tool (**Figure 3B-D**). We identified and quantified between 25.000 to 27.000 peptides derived from 4800 to 5000 proteins in each of the individual *E*.*coli*:HEK samples (**Table 2**). Moreover, the coefficients of variation (CV) of the signal for the different transitions per condition across the replicates (n = 4) are similar for the different *E*.*coli*:HEK ratios (**Figure 4A**). Differential expression analysis using the MSstats tool revealed significantly dysregulated proteins between the different *E*.*coli*:HEK ratios. As expected when comparing the 1:17 against the 1:7 *E*.*coli*:HEK ratio, multiple *E*.*coli* proteins are significantly downregulated in the 1:17 *E*.*coli*:HEK samples (**Figure 4B**). Furthermore, some human proteins appear to be upregulated in this comparison, which might be due to displacement effects by the added amount of *E*.*coli* proteome. Even when comparing the two lowest *E*.*coli*:HEK ratios (1:50 vs 1:17) significantly dysregulated *E*.*coli* proteins can be detected, highlighting the overall functionality as well as a suitable sensitivity of the DIA analysis pipeline in Galaxy (**Supplementary Figure 6**). The complete analysis was performed on the European Galaxy instance ^22^. All analysis histories, as well as workflows, are available in the supporting information of this publication, to ensure full reproducibility of the presented results. Accurate, transparent and complete sharing of parameters and whole analysis is greatly simplified by Galaxy’s intrinsic features for sharing and publishing. Shared histories and workflows as well as the Galaxy software fulfill the FAIR principles ^24^.

**Table 2.** Overview of identified and quantified precursors, peptides and proteins.

| *E.coli*:HEK ratio | Replicate | Precursors | Peptides | Proteins |
| --- | --- | --- | --- | --- |
| 1:17 | 1 | 28792 | 27117 | 4979 |
| 1:17 | 2 | 28697 | 27040 | 4972 |
| 1:17 | 3 | 28673 | 27024 | 4970 |
| 1:17 | 4 | 28641 | 27003 | 4989 |
| 1:3 | 1 | 28711 | 27060 | 5011 |
| 1:3 | 2 | 28690 | 27043 | 5000 |
| 1:3 | 3 | 28672 | 27028 | 5006 |
| 1:3 | 4 | 28669 | 27035 | 4996 |
| 1:50 | 1 | 28191 | 26576 | 4906 |
| 1:50 | 2 | 28255 | 26636 | 4914 |
| 1:50 | 3 | 28231 | 26595 | 4919 |
| 1:50 | 4 | 28192 | 26577 | 4916 |
| 1:7 | 1 | 28833 | 27160 | 4994 |
| 1:7 | 2 | 28791 | 27123 | 5006 |
| 1:7 | 3 | 28804 | 27136 | 5004 |
| 1:7 | 4 | 28837 | 27166 | 5011 |
| HEK only | 1 | 27166 | 25669 | 4843 |
| HEK only | 2 | 27182 | 25683 | 4862 |
| HEK only | 3 | 27176 | 25663 | 4869 |
| HEK only | 4 | 27099 | 25600 | 4858 |

DIA analysis results were filtered at 1% FDR on peptide and protein level during export using the PyProphet export tool. In combination with a sample annotation file the swath2stats functionality was applied yielding an overview of identified and quantified precursors, peptides and proteins in each sample.

**Figure 4.**
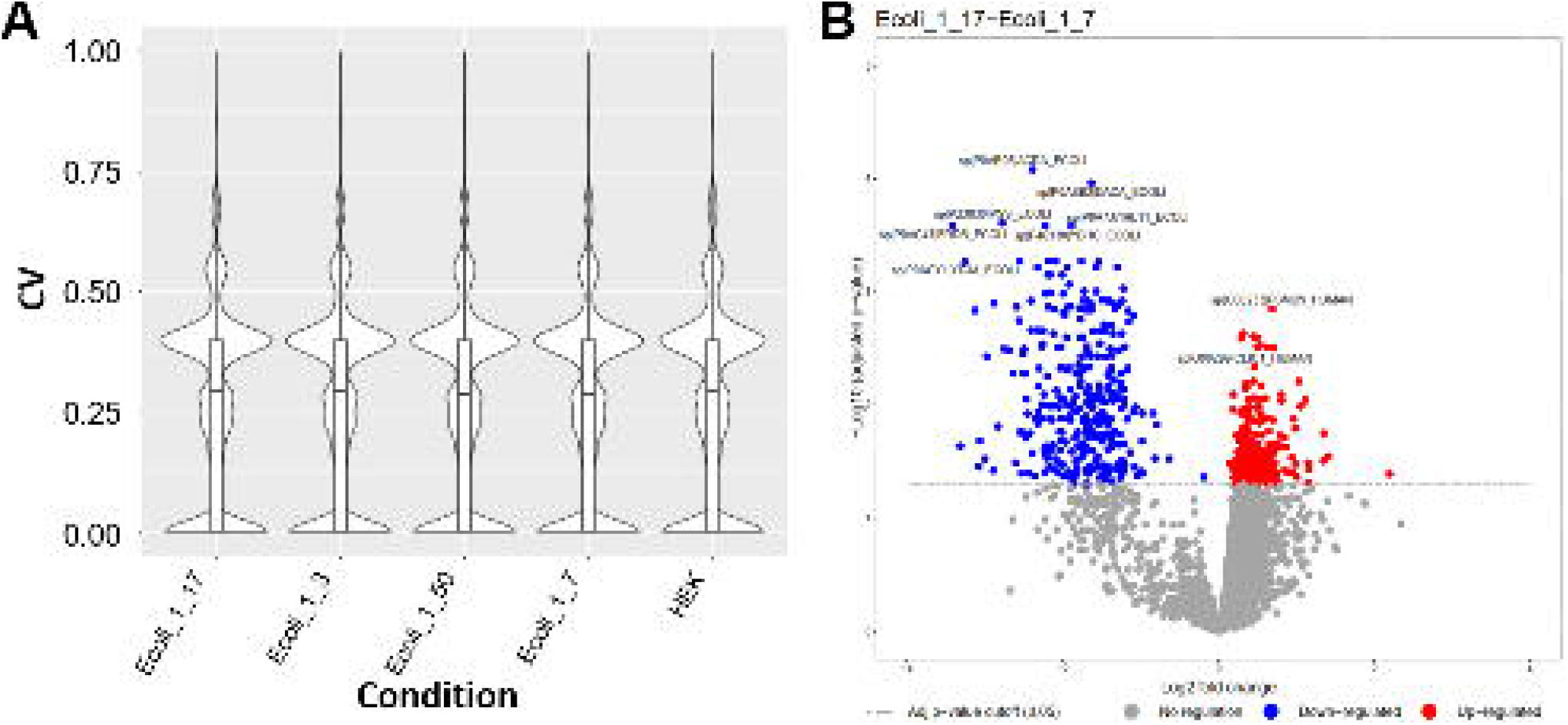
Results obtained using the DIA analysis tools in Galaxy. (A) Violin plot illustrating the distribution of coefficients of variation (CV) of the signal for the different transitions per condition (here *E*.*coli*:HEK ratios) across replicates (each n = 4). (B) Volcano plot showing -log10 adjusted p-values against log2 fold changes, highlighting differentially expressed proteins comparing the two *E*.*coli*:HEK ratios 1:17 versus 1:7.

## Conclusions

To conclude, our DIA Galaxy tools and workflows represent a powerful and user friendly software solution for the analysis of large-scale DIA experiments. We developed DIA analysis tools based on open-source software such as OpenSwath, PyProphet, diapysef and swath2stats, that can be integrated with existing tools providing a flexible and modular analysis pipeline. Moreover, the tools can be assembled into complete DIA analysis workflows promoting straightforward and reproducible analysis of large sample cohorts. All tools are accessible via the Galaxy system of graphical user interfaces and have access to public clouds. The web-based access in Galaxy and the extensive training material empower a broad community of researchers to perform their DIA analysis, without the need for enhanced computational skills and resources. Complete analysis histories and workflows can be shared and published via Galaxy promoting transparent and reproducible DIA analysis. By integrating a suite of modular DIA tools in Galaxy and presenting fit-for-purpose, readily usable DIA workflows, we make the abilities and reproducibility of the Galaxy infrastructure accessible to the DIA proteomics community.

## Availability of Supporting Source Code and Requirements

Project name: Data-independent acquisition analysis workbench in Galaxy

RRID number: SCR_017410 (https://scicrunch.org/resolver/RRID:SCR_017410)

Project homepage: https://github.com/galaxyproteomics/tools-galaxyp

Galaxy Toolshed: https://toolshed.g2.bx.psu.edu/

Operating system(s): Unix

Training repository: https://galaxyproject.github.io/training-material/proteomics

License: MIT

## Availability of Supporting Data and Materials

Galaxy workflow to generate a spectral library (https://usegalaxy.eu/u/matthiasfahrner/w/dia-lib-hek-ecoli-3eg-data);

Galaxy workflow to perform DIA analysis (https://usegalaxy.eu/u/matthiasfahrner/w/dia-analysis-using-hek-ecoli-3-eg-data);

Galaxy workflow to perform the statistical analysis:

a) using swath2stats converted MSstats input (https://usegalaxy.eu/u/matthiasfahrner/w/hek-ecoli-dia-statistics-swath2stats-3eg-data),

b) using pyprophet export tsv (https://usegalaxy.eu/u/matthiasfahrner/w/hek-ecoli-dia-statistics-3eg-data-1);

Galaxy history of the spectral library generation (https://usegalaxy.eu/u/matthiasfahrner/h/dia-lib-hek-ecoli-3eg-data);

Galaxy history of the DIA analysis including statistical analysis (https://usegalaxy.eu/u/matthiasfahrner/h/dia-analysis-statistics-hek-ecoli-3eg-data).

Mass spectrometry data has been deposited and is available via MassIVE (Username:MSV000087859_reviewer ; Password: DIA_in_Galaxy).

## Additional Files

**Supplementary Figure 1. Galaxy workflows for statistical analysis of DIA data**. Galaxy workflow for statistical analysis of DIA data with MSstats using a group comparison matrix file and (A) the swath2stats processed PyProphet export results or (B) the direct PyProphet export results and a sample annotation file.

**Supplementary Figure 2. Overview of Target-Decoy scoring results using PyProphet peptide with experiment-wide peptide-level error-rate control**.

(A) Receiver operating characteristic (ROC) curve highlighting the sensitivity and specificity of the Target-decoy scoring. (B) Blot showing the discriminatory score (d-score) performance between target (green) and decoy (red) transitions. (C) and (D) Barblot and density plot showing d-score distribution among target (green) and decoy (red) transitions. (E) Histogram showing the distribution of p-values computed based on the target-decoy scoring.

**Supplementary Figure 3. Overview of Target-Decoy scoring results using PyProphet peptide with global peptide-level error-rate control**

(A) Receiver operating characteristic (ROC) curve highlighting the sensitivity and specificity of the Target-decoy scoring. (B) Blot showing the discriminatory score (d-score) performance between target (green) and decoy (red) transitions. (C) and (D) Barblot and density plot showing d-score distribution among target (green) and decoy (red) transitions. (E) Histogram showing the distribution of p-values computed based on the target-decoy scoring.

**Supplementary Figure 4. Overview of Target-Decoy scoring results using PyProphet protein with experiment-wide protein-level error-rate control**.

(A) Receiver operating characteristic (ROC) curve highlighting the sensitivity and specificity of the Target-decoy scoring. (B) Blot showing the discriminatory score (d-score) performance between target (green) and decoy (red) transitions. (C) and (D) Barblot and density plot showing d-score distribution among target (green) and decoy (red) transitions. (E) Histogram showing the distribution of p-values computed based on the target-decoy scoring.

**Supplementary Figure 5. Overview of Target-Decoy scoring results using PyProphet protein with global protein-level error-rate control**.

(A) Receiver operating characteristic (ROC) curve highlighting the sensitivity and specificity of the Target-decoy scoring. (B) Blot showing the discriminatory score (d-score) performance between target (green) and decoy (red) transitions. (C) and (D) Barblot and density plot showing d-score distribution among target (green) and decoy (red) transitions. (E) Histogram showing the distribution of p-values computed based on the target-decoy scoring.

**Supplementary Figure 6. Results obtained using the DIA analysis tools in Galaxy**. Volcano plot showing -log10 adjusted p-values against log2 fold changes, highlighting differentially expressed proteins comparing the two *E*.*coli*:HEK ratios 1:50 versus 1:17.

### Abbreviations

CV: Coefficients of variation
DDA: Data-dependent acquisition
DIA: Data-independent acquisition
FAIR: findable, accessible, interoperable, and re-usable
FDR: False discovery rate
LC-MS/MS: Liquid chromatography-tandem mass spectrometry
MIAPE: minimum information about a proteomics experiment
OSW: OpenSwathWorkflow
PQP: peptide query parameter
RT: retention time
TSV: tab-separated values.

## Authors Contribution

M.F. conceived the project, tested the tools, and developed the training material and the case study. M.C.F. developed the Galaxy tool wrappers and the training material, and contributed to the conceptualization. B.A.G. and M.B. developed the Galaxy tool wrapper.

H.R. developed parts of the applied software. B.A.G and O.S. contributed to the conceptualization, methodology and funding acquisition. M.F. wrote the manuscript. All authors critically read and approved the manuscript’s contents.

## Acknowledgments

The authors acknowledge the support of the Freiburg Galaxy Team, Bioinformatics, University of Freiburg (Germany) funded by the <u>Collaborative Research Centre 992 Medical Epigenetics</u> (<u>DFG</u> grant SFB 992/1 2012) and the German Federal Ministry of Education and Research <u>BMBF</u> grant 031 A538A <u>de.NBI</u>-RBC.

