## Supplementary figures and images for "Democratizing Data-Independent Acquisition Proteomics Analysis on Public Cloud Infrastructures Via The Galaxy Framework"

### Supplementary Figure 1

#### Slide 1
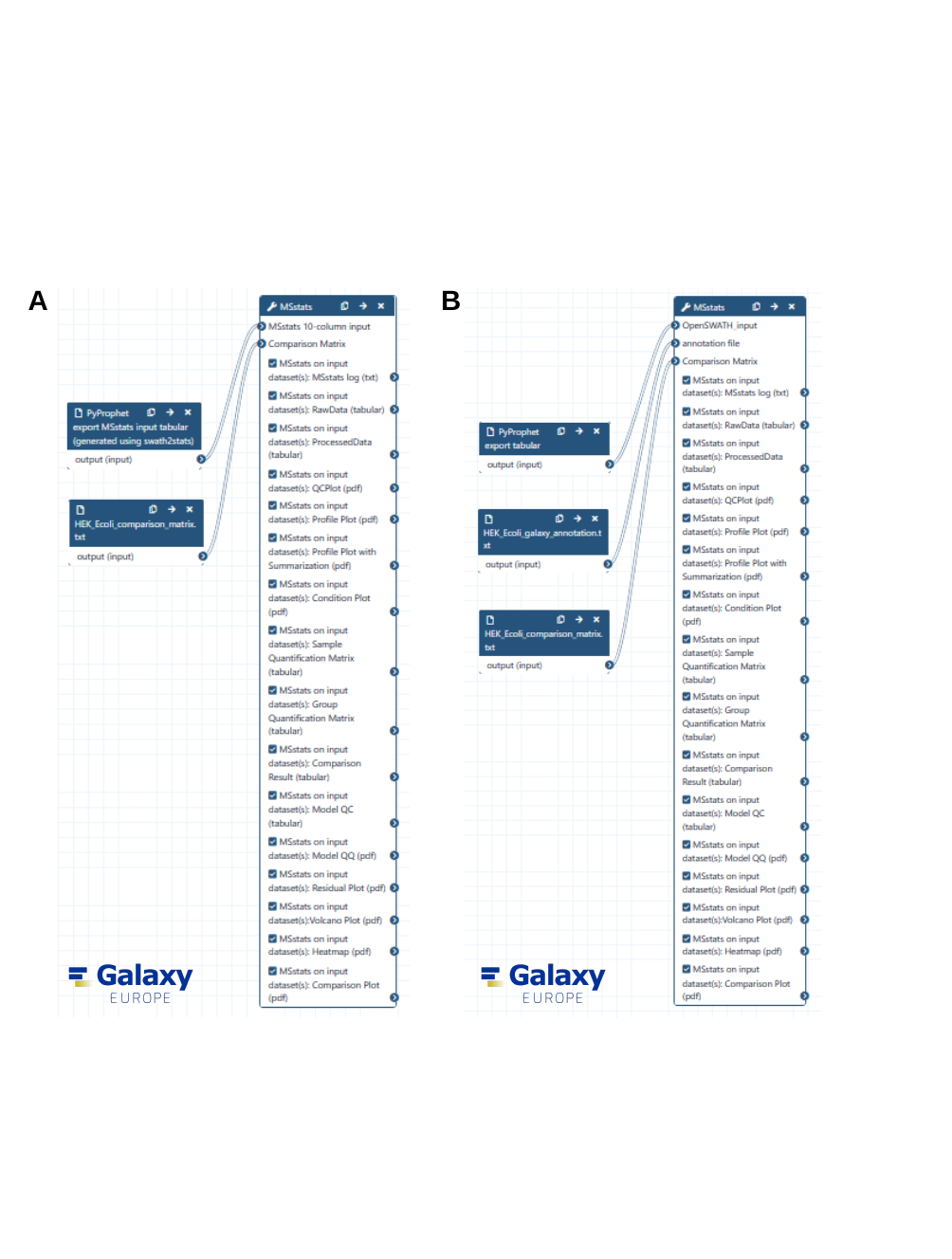

A
B

### Supplementary Figure 2

#### Slide 1
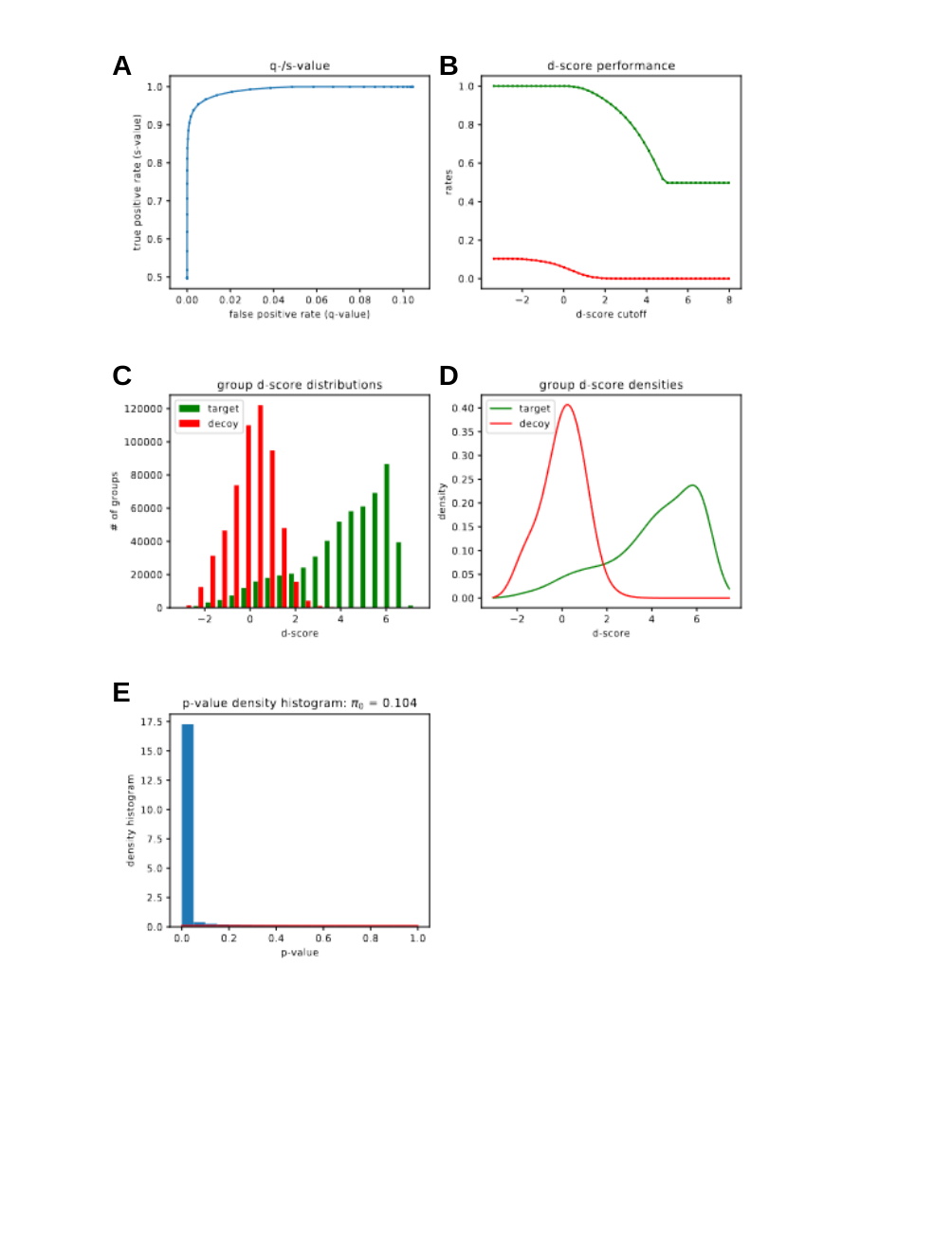

A
B
C
D
E

### Supplementary Figure 3

#### Slide 1
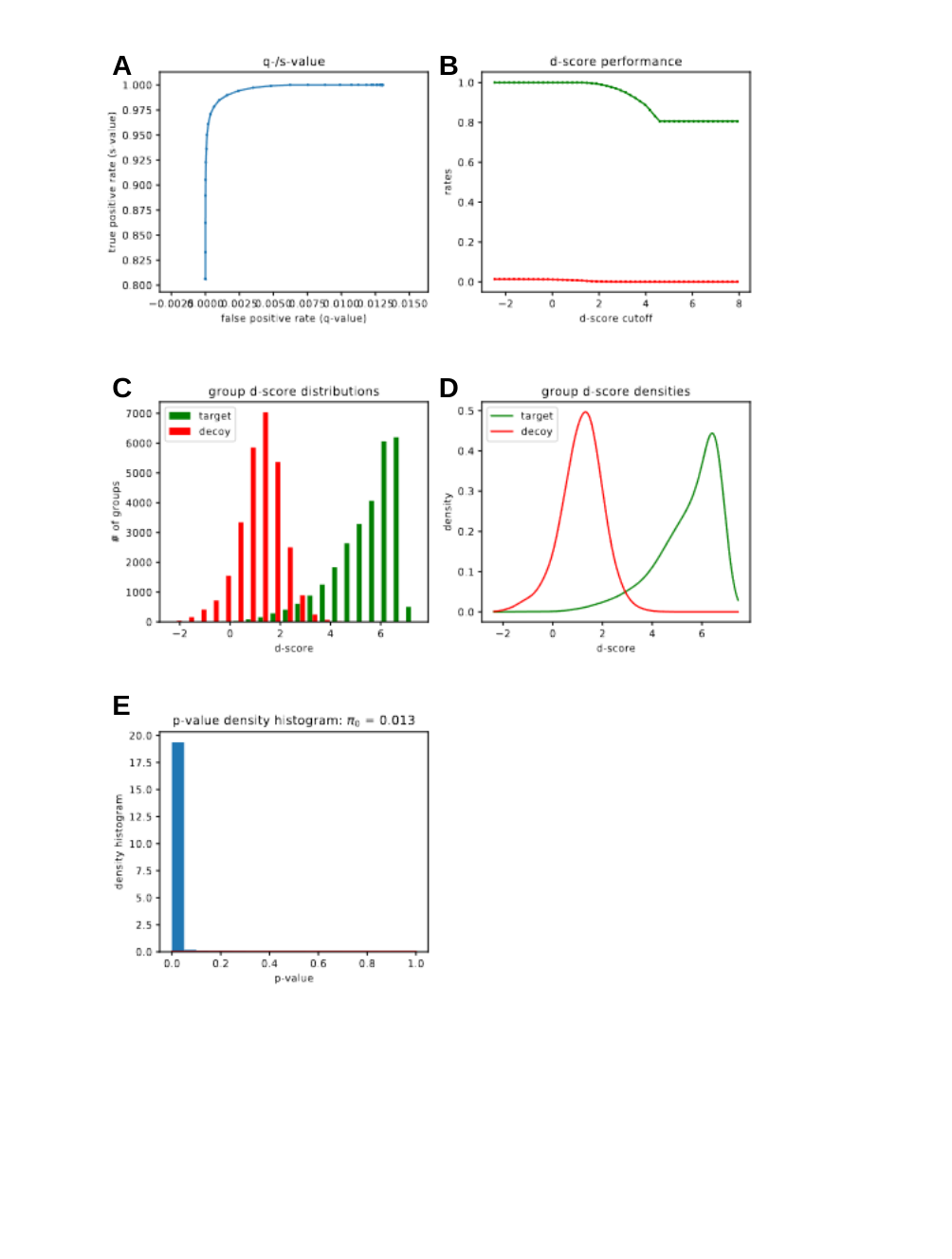

A
B
C
D
E

### Supplementary Figure 4

#### Slide 1
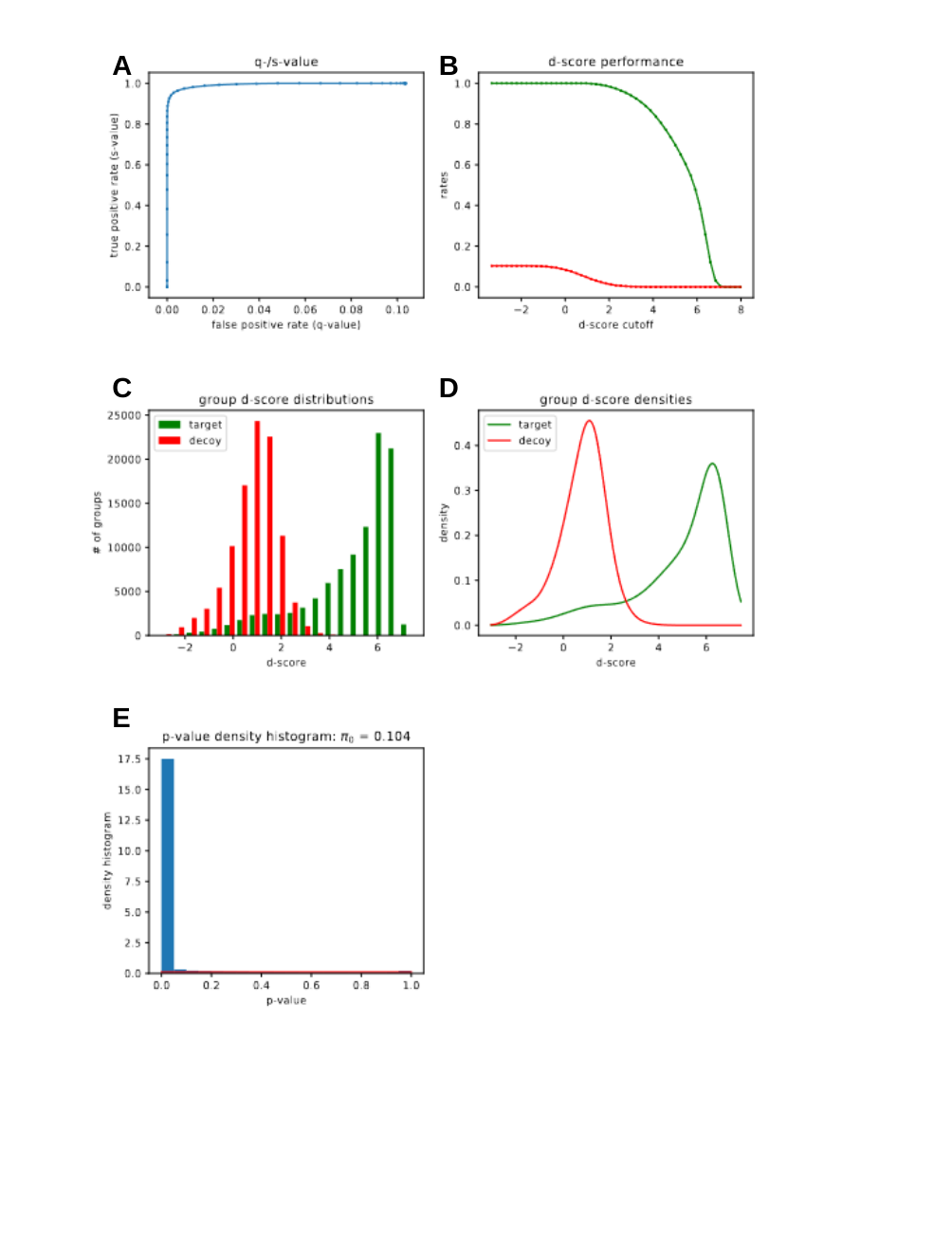

A
B
C
D
E

### Supplementary Figure 5

#### Slide 1
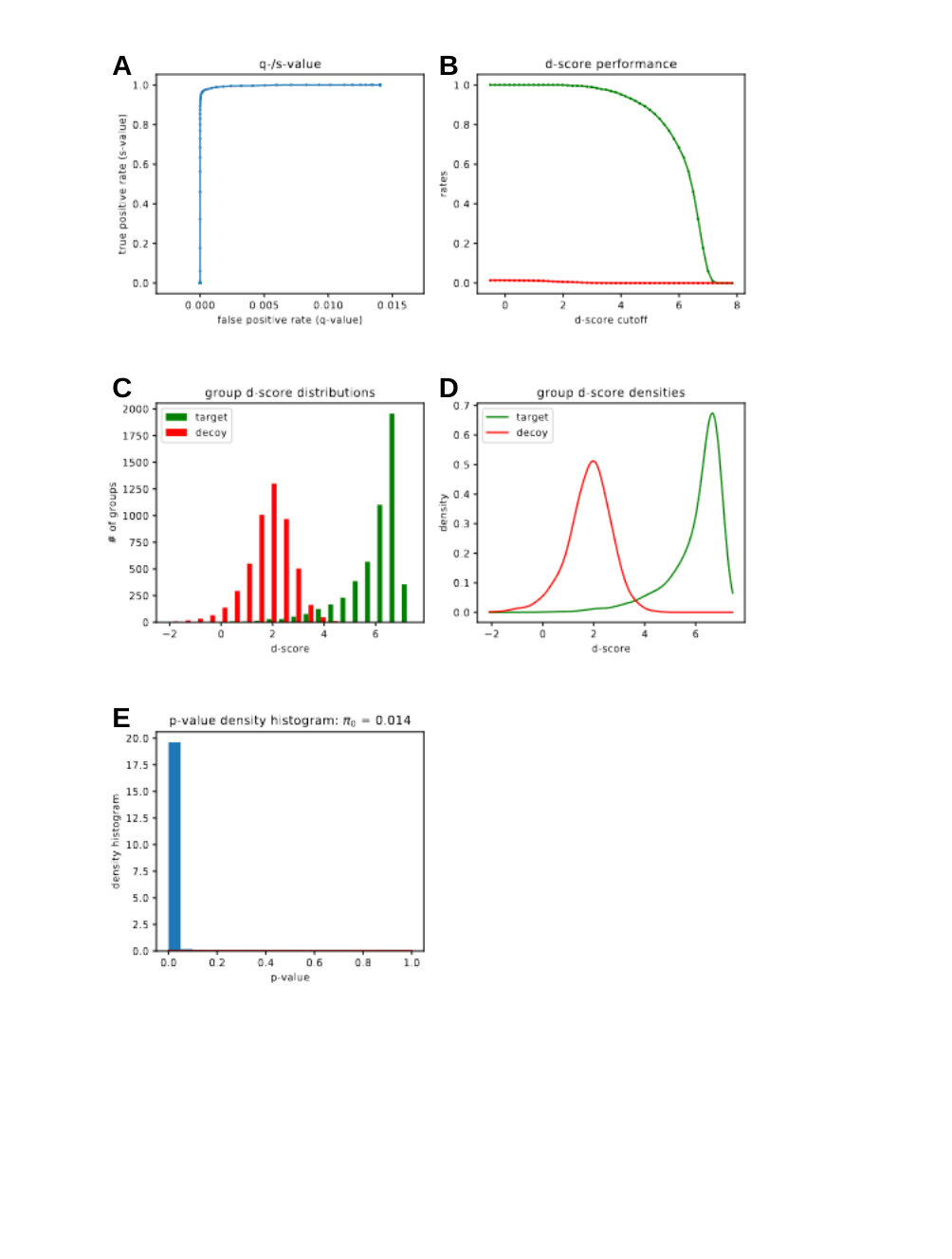

A
B
C
D
E

### Supplementary Figure 6

#### Slide 1
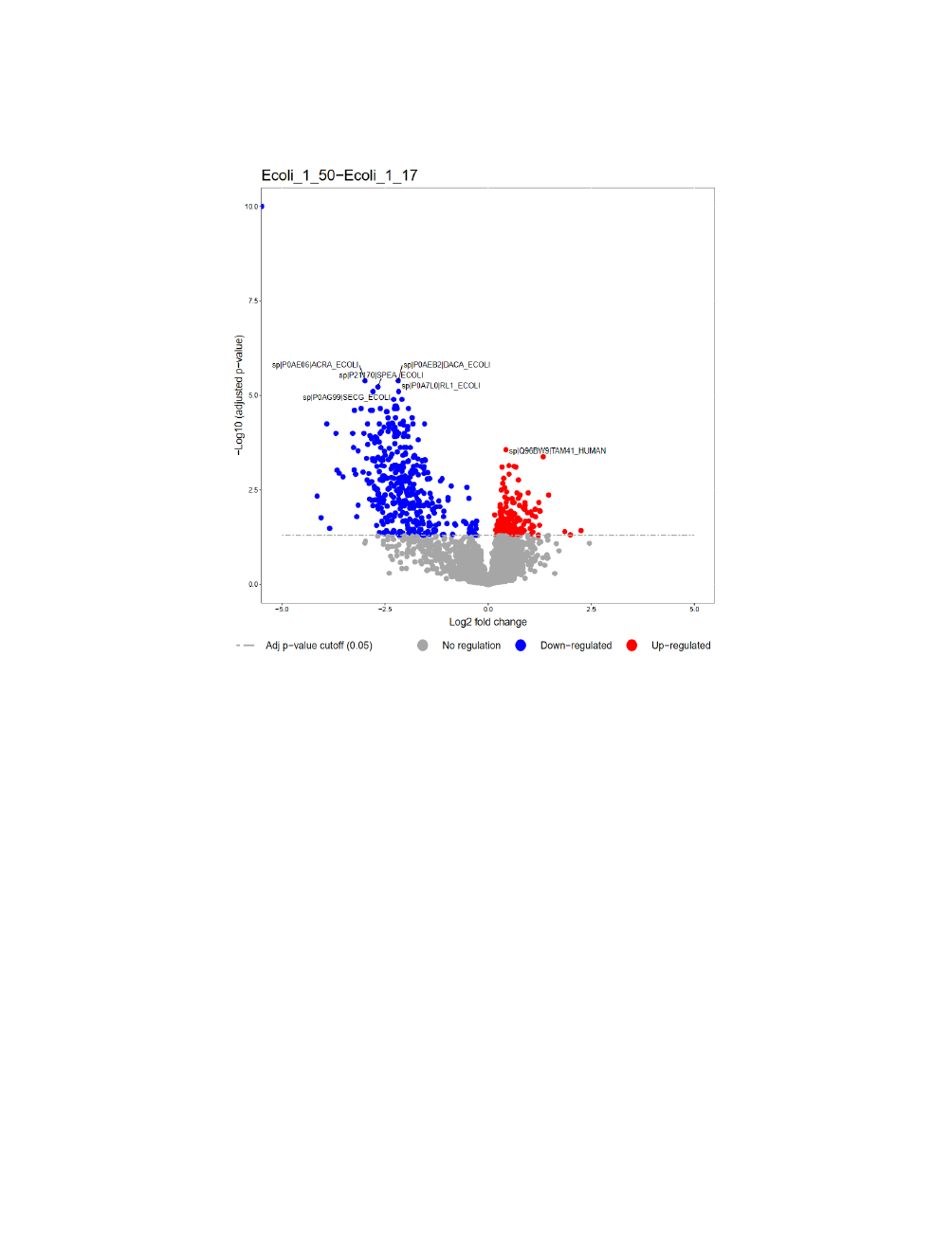
